# Nanotopography boosts cellular uptake by inducing macropinocytosis

**DOI:** 10.1101/2023.12.20.571698

**Authors:** Morteza Aramesh, Di Yu, Magnus Essand, Cecilia Persson

## Abstract

Efficient cellular uptake of biomolecules, including genetic material, mRNA, proteins, and nanoparticles, requires novel approaches to overcome inherent cellular barriers. This study investigates how nanotopographical cues from nanoporous surfaces impact the uptake efficiency of diverse molecules by cells. The results demonstrate that cellular uptake efficiency increases significantly on nanoporous surfaces compared to flat surfaces. Notably, this process is found to be dependent on the size and morphology of the nanopores, reaching its peak efficacy with blind pores of 400 nm in size. Enhanced genetic transduction on nanoporous surfaces were observed for multiple vectors, including lentiviruses, baculoviruses, and mRNA molecules. The versatile nature of this approach allows co-transfection of cells with multiple mRNA vectors. Moreover, the nanoporous platform was used for efficient and fast manufacturing of CAR-T cells through lentiviral transduction. Furthermore, we pinpoint macropinocytosis as the predominant mechanism driving increased cellular uptake induced by the nanoporous surfaces. The method introduced here for enhancing genetic transduction of cells has applications in immunotherapy research, drug delivery, and cell engineering.

## Introduction

Efficient cellular uptake of genetic material like DNA or RNA, proteins, and nanoparticles is crucial for emerging gene therapies for treating genetic disorders and cancers.^1,2^ Cell-based therapies, such as self-renewing hematopoietic stem cells and T cells for immunotherapy, are on the rise in clinical trials.^3,4^ Despite their significance, efficient delivery of genetic material to stem cells and immune cells remains a challenge, due to low transduction efficiency and compromised cellular fitness after transduction.^5,6^

Various strategies are employed to improve successful gene deliveries.^2,7^ These include mechanical techniques such as electroporation, chemical approaches involving lipid or nanoparticle carriers, and biologically assisted methods using viral or bacterial vectors.^8–11^ Currently, viral vectors are the clinical standard due to their relatively higher efficiency and precision.^12^ A notable example includes the utilization of lentiviral transduction in cancer immunotherapy for Chimeric Antigen Receptor (CAR) T cell therapy, where CAR-T cell production remains suboptimal due to the low transduction efficiency of T cells, highlighting the demand for technologies to improve transfection of immune cells.^4,12^ Moreover, transduction with multiple vectors proves significantly less efficient compared to single vectors, thereby restricting the potential applications of cell engineering that involve targeting multiple genes.^13^

The application of cell-instructive materials to guide cellular responses, especially in delivery systems, represents a state-of-the-art approach. Typically designed at the nanoscale or microscale, these materials provide cues to cells through biochemical signals and physical attributes like stiffness and topography.^14–19^ They have been engineered to guide cellular activities such as metabolism, proliferation, differentiation, secretion, and cellular uptake.^20^

Here, we demonstrate that nanoporous materials can act as cell-instructive materials to guide cellular uptake by enhancing delivery of lentiviruses, baculoviruses, mRNA, antibodies, and liposomes, as measured by transduction efficiency. We show the efficacy of this process is influenced by the size and shape of the nanopores. Moreover, we highlight practical applications in production of CAR-T cells and multiple mRNA constructs delivery. Additionally, through inhibition of individual endocytic pathways, we identify macropinocytosis as the primary mechanism responsible for enhanced cellular uptake in cells induced by nanoporosity of the surface.

## Results and Discussion

### Enhanced uptake on nanoporous surfaces

Figure 1a illustrates the schematic representation of the workflow employed to investigate cellular uptake on nanoporous surfaces. The nanoporous surfaces were primarily composed of polycaprolactone (PCL), unless otherwise specified. These surfaces featured an average pore size ranging from 100 nm to 1000 nm. Non-porous PCL films were used as control samples. Cells were directly cultured on these films located at the base of standard well-plates. The initial interaction of the cells with the nanoporous surface was observed using confocal microscopy (Figure 1c**,d**). The distinctive dotted structure under the microscope is an indication of formation of actin-rich protrusions into the nanopores. We have previously shown that these protrusions occur at pore sizes larger than 100 nm.^16^ The cross-section profile of a cell on a 400 nm porous surface is shown in Figure 1c, where the extent of the penetration into the depth of the pores is clearly visible.

**Figure 1.**
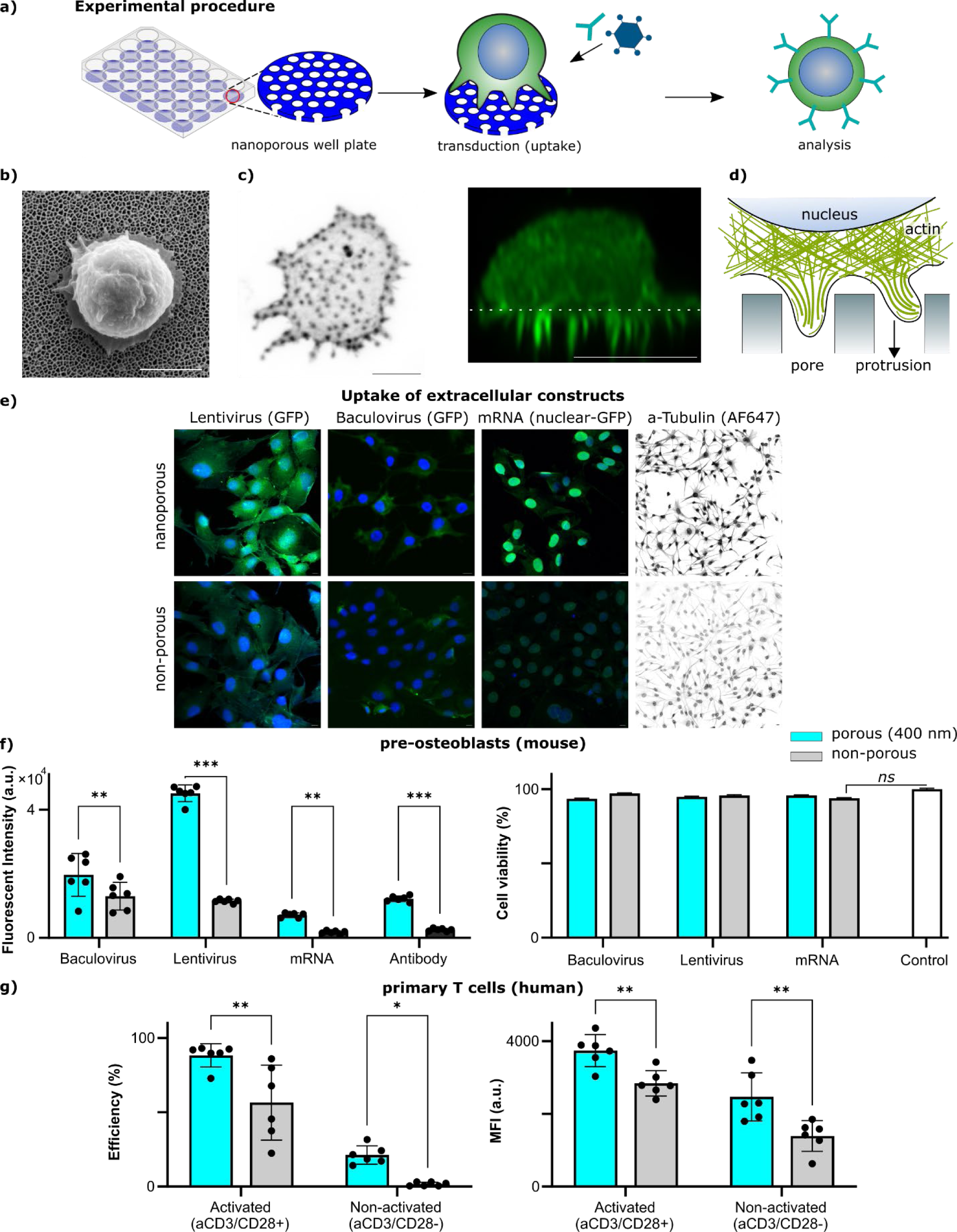
**a)** A schematic illustration of the experimental procedure, depicting the coating of culture plates with nanoporous films, usually 400 nm in average pore size for optimal extracellular uptake efficiency (not to scale). **b)** Scanning electron microscope image of a primary human T cell on top of a porous film (scale bar, 5 μm). **c)** Confocal fluorescence microscopy images of actin cytoskeleton of primary human cells on a porous film with 400 nm pore size. (left) a top view and (right) a cross-section view (scale bar, 5 μm). **d)** A schematic presentation of the actin-rich protrusions in T cells on a porous surface (not to scale). **e)** Fluorescence microscopy images of (mouse) pre-osteoblast cells cultured on porous (400 nm) and non-porous PCL films and the uptake of different vectors: GFP-lentivirus, GFP-baculovirus, nuclear-GFP mRNA and anti-tubulin antibody (AF647). The nucleus was stained with Hoechst. **f)** Quantitative measurement of uptake via optical measurements using a plate reader. **g)** Flow cytometry analysis of primary human T cells transfected with GFP-lentivirus on porous (400 nm) and non-porous PCL films. aCD3/CD28± indicates the presence or absence of activating antibodies prior to transfection (24 hours). The efficiency was assessed by determining the relative number of GFP+ cells in relation to the total cell count. MFI is the mean fluorescence intensity of GFP in GFP+ cells. A two-way ANOVA test was conducted to assess the statistical significance (*: p < 0.05, **: p < 0.01, ***: p < 0.001).

Two types of cells were employed for quantifying cellular uptake: mouse pre-osteoblast cells (adherent) and primary human T cells (non-adherent). Genetic materials or antibodies were introduced into the cell culture medium 24 hours after seeding the cells on the surfaces, without the use of additional chemical reagents, additives, or enhancers. The genetic constructs tested included GFP-lentivirus, GFP-baculovirus, and nuclear-GFP mRNA. To assess antibody uptake, anti-tubulin tagged with a fluorescent dye (AF647) was employed. The cells were incubated with the genetic constructs or antibodies for 24 hours before subsequent measurements. The success of uptake or expression, as well as cell viability, was evaluated. For adherent cells, fluorescent intensity of transduced cells was determined using a plate reader, and live/dead staining was employed to assess cell viability for each construct. In the case of non-adherent cells, flow cytometry was utilized to quantify the efficiency and strength of transduction achieved by GFP encoding lentiviruses. Transduction efficiency was evaluated by measuring the percentage of cells expressing the respective biomolecule construct after exposure to porous and non-porous surfaces.

For adherent cells (mouse pre-osteoblasts), the results showed that the fluorescent intensity of transduced cells on porous surfaces (400 nm) was significantly higher compared to non-porous surfaces when using baculovirus, lentivirus, mRNA, or antibodies (Figure 1e**,f**). Additionally, live/dead staining indicated high cell viability for all constructs on both surface types (Figure 1f). In the case of non-adherent cells (primary human T cells), flow cytometry demonstrated that transduction efficiency is much higher on porous surfaces (400 nm) compared to non-porous surfaces. Transduction was performed using GFP lentiviruses for both activated (αCD3/CD28+) and non-activated T cells (αCD3/CD28-) (Figure 1g). Both activated and non-activated T cells exhibited higher transduction efficiency on porous surfaces. Remarkably, activated T cells exhibited a transduction efficiency of 98% on porous surfaces. The higher transduction rate was accompanied by increased GFP expression at the individual cell level, measured by the mean fluorescent intensity (MFI). The 4-fold increased transduction rate of activated T cells compared to non-activated cells can be attributed to the heightened metabolic activity, increased surface area (volume), and active proliferation of activated cells, factors that contribute to facilitating the entry of genetic material into the nucleus.

In summary, these findings suggest that porous surfaces (400 nm) enhance cellular uptake, integration of genetic material, and, consequently, the transduction efficiency for various constructs and diverse cell types.

### Uptake Characterization

We explored surfaces featuring a range of pore sizes, from 100 nm to 1000 nm, and observed enhanced cellular uptake across ranges of pore diameters (Figure 2). A substantial increase in uptake efficiency was evident on surfaces with 400 nm pores, using nuclear-GFP mRNA, whereas nuclear transduction was notably less effective on non-porous surfaces (Figure 2a**,b**). The cytosolic leakage of nuclear GFP in cells cultured on substrates with 400 nm pores may be attributed to protein overexpression, which was intensified after 48 hours (**Figure S1**). There was no obvious liners correlation between cellular uptake and pore sizes of 100-1000 nm, and the 400 nm pores led to the highest uptake, suggesting a pore-size dependent mechanism which makes it distinct from the uptake mechanism on non-porous surfaces.

**Figure 2.**
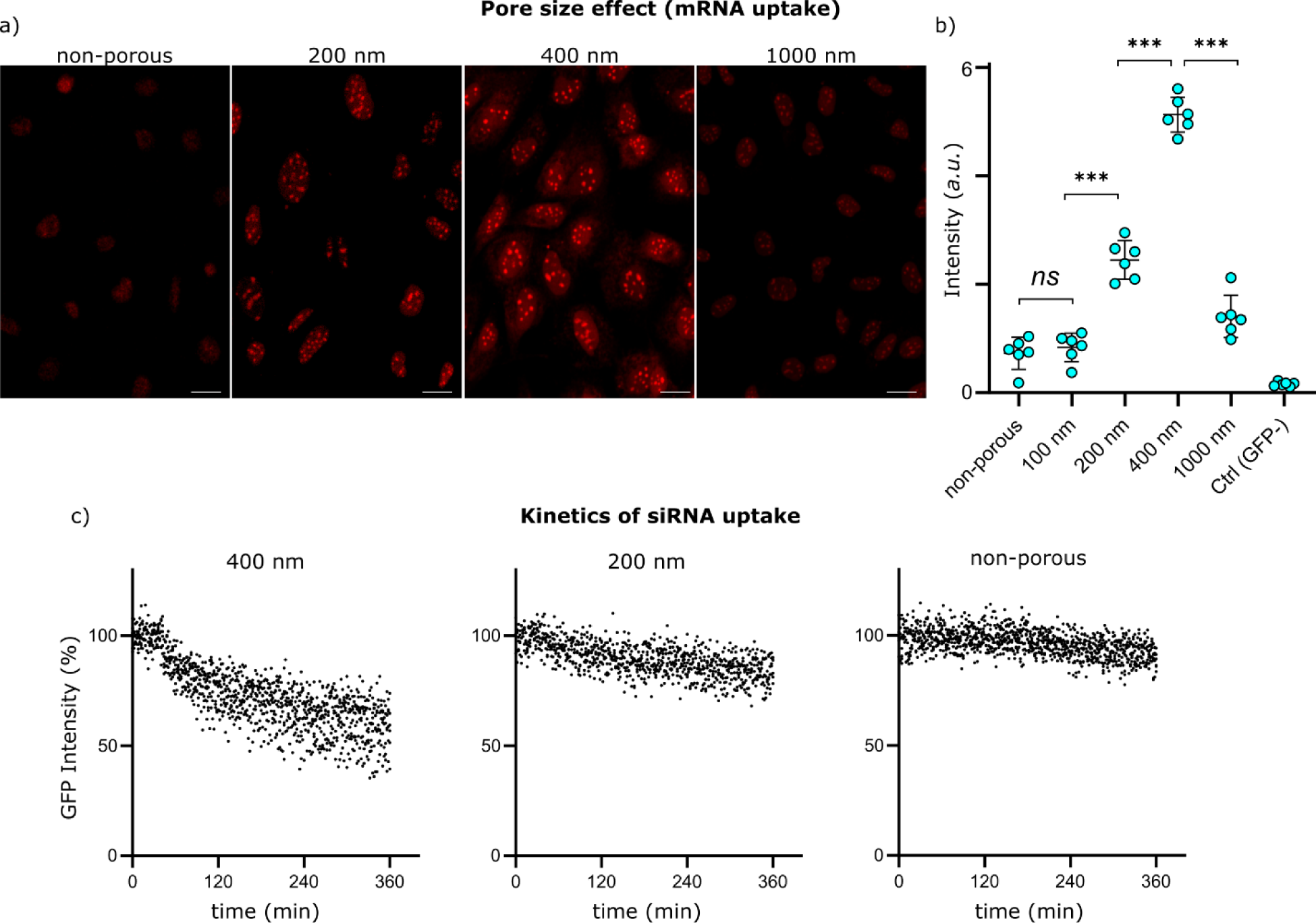
Pore size effect. **a)** Confocal fluorescence microscopy images of mouse pre-osteoblast cells transduced with nuclear-GFP mRNA upon seeding on PCL films with varied pore diameters, including non-porous, 200, 400, and 1000 nm (scale bar, 5 μm). **b)** Quantitative uptake measurements were conducted using optical readings with a plate reader. A scrambled mRNA (GFP-) served as the control. Statistical significance was evaluated through a two-way ANOVA test (*: p < 0.05, **: p < 0.01, ***: p < 0.001). **c)** Kinetics analysis of siRNA uptake by mouse pre-osteoblast cells on 400 nm, 200 nm, and non-porous films. The decline in GFP signal was an indicator of siRNA uptake and subsequent endosomal escape into the cytosol.

Moreover, using porous materials with different pore shapes showed that pore geometry has an impact on enhancing the transduction process. A significant difference was observed in transduction efficiency between blind pores and open pores with straight channel walls (**Figure S2**). The blind pores exhibited a higher effectiveness in transducing cells compared to the straight-walled pores. The depth of the blind pores is within the order of the pore diameter, while straight-walled pores were approximately 100 μm deep (**Figure S3**).

The use of chemical enhancers, such as polybrene, has shown to enhance the transduction efficiency, particularly through charge neutralization.^21^ Polybrene is a positively charged polymer that interacts with the cell membrane (negatively charged), facilitating the entry of genetic material into cells which are also usually negatively charged (such as mRNA, DNA and viruses). The application of polybrene yielded a significant increase in cell transduction on non-porous surfaces, however, its influence was less pronounced on porous surfaces (**Figure S4**). These results also point to a potentially distinct mechanism of uptake on porous vs non-porous surfaces, as the uptake on porous surfaces is not affected by electrostatic charge neutralization.

To gain insights into the dynamics of cellular uptake, we performed a comparative analysis by examining the decay of fluorescent intensity in GFP-expressing cells with a GFP quencher using small interfering RNA (siRNA). These analyses were conducted on both porous and non-porous surfaces at 37 °C. The reduction in GFP signal served as an indicator of siRNA uptake and subsequent endosomal escape into the cytosol. A significantly faster decrease in GFP signal on surfaces with 400 nm pores was observed compared to those with 200 nm pores and non-porous surfaces, indicated by the steeper decay of GFP intensity over time (Figure 2c). On surfaces with 400 nm pores, GFP intensity was reduced to 60% within the 6 hour measurement, and the decay rate reached its maximum within 60 minutes (**Figure S5**). On the other hand, GFP intensity was reduced to 85% and 95% on 200 nm pores and non-porous surfaces, respectively. Also the decay rate increased gradually over time and with a 3.5 times slower rate compared to 400 nm surfaces (**Figure S5**).

Furthermore, we investigated the impact of temperature on cellular uptake. When cells were exposed to siRNA at 4 °C for durations of 30 minutes, 1 hour, 2 hours, and 4 hours, the GFP signal remained largely unchanged both on porous (400 nm) and non-porous surfaces (**Figure S6**).

### Applications

We explored the use of enhanced delivery methods with porous materials for two demanding biological processes: i) co-transfection with multiple mRNA vectors and ii) the manufacturing of CAR-T cells. These procedures are known for their complexities and conventionally low efficiencies. Here we compare the effectiveness of our approach with conventional methods in addressing these challenges.^12,22,23^

#### Co-transfection

Co-transfection involves introducing multiple vectors, such as mRNA or plasmids, into host cells concurrently. Achieving efficient co-transfection presents challenges, primarily due to the inherent probability factors associated with cellular entry, endosomal escape and expression processes. Therefore, enhanced cellular uptake can increase the likelihood of successful multiple vector transfection.

We sought to investigate the efficiency of gene transfection on both non-porous and porous surfaces (400 nm) and examine the probabilities of achieving single, double, and triple transfections with three distinct mRNA constructs. mRNA constructs were encapsulated within lipofectamine lipid nanoparticles, and co-transfection was performed on T cells immediately after seeding on the surfaces. After 24 hours, we assessed the transfection efficiency using flow cytometry and analyzed the expression levels of the proteins in the transfected cells (Figure 3). The results indicate decaying probabilities for single, double, and triple transfections on non-porous surfaces. Porous surfaces (400 nm), however, substantially increased the likelihood of transfection, with double vector transfection showing the highest probability. Approximately 50% of the cells exhibited expression from at least two vectors. Approximately 10% of the cells showed expression from the three vectors, compared to <1% on non-porous surfaces, which is an order of magnitude enhancement in the successful uptake. Additionally, the fluorescent intensity, indicative of a higher level of expression per cell, was significantly greater on porous surfaces compared to non-porous surfaces (Figure 3b). These results underscore the potency of nanoporous materials for multiple-vector cellular transfection.

**Figure 3.**
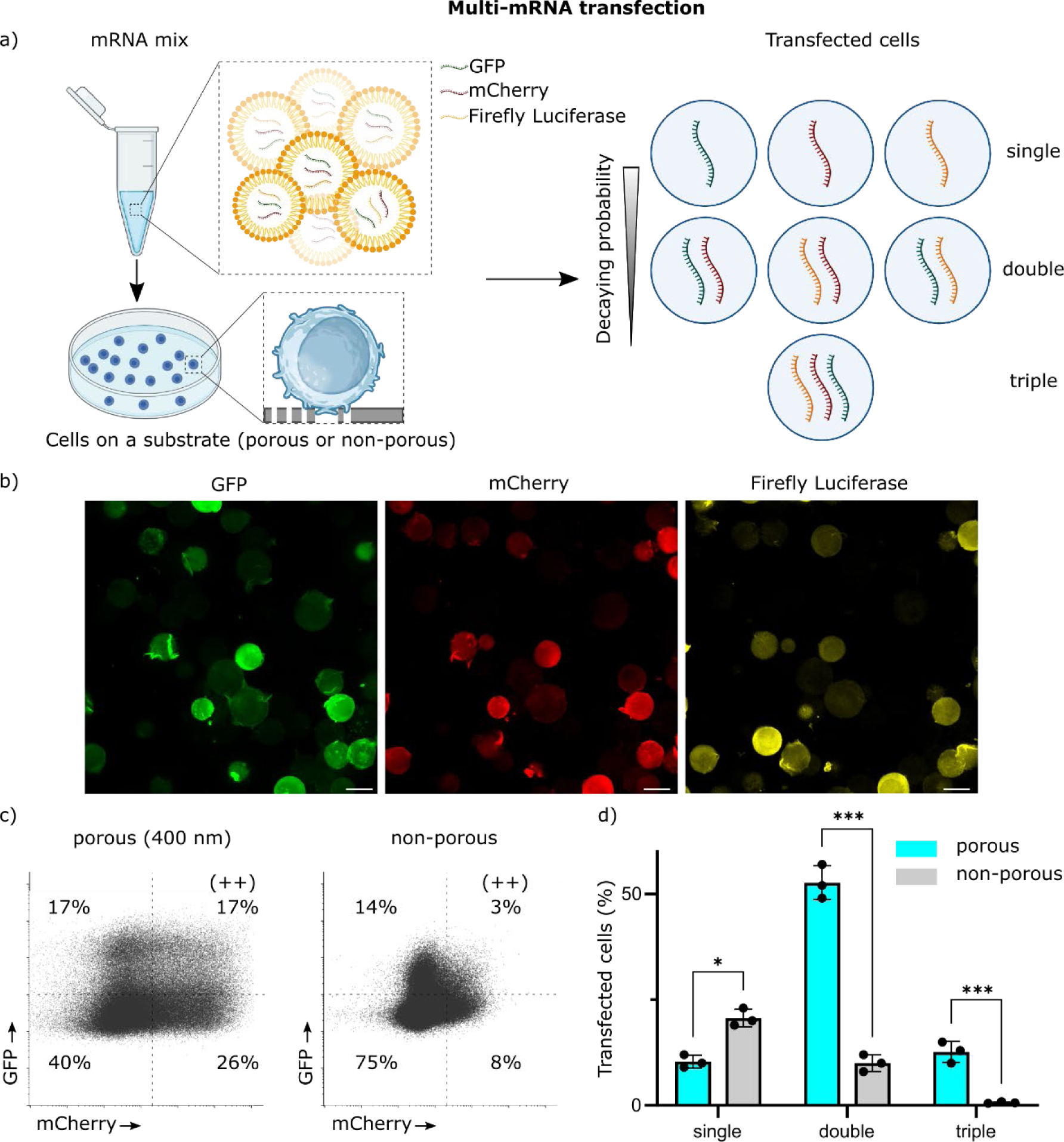
Multiple mRNA uptake. **a)** A schematic depiction of the experimental procedure: three distinct mRNA constructs were combined and enclosed within lipid nanoparticles. These nanoparticles were subsequently introduced to T cells cultivated on either porous (400 nm) or non-porous surfaces. Typically, achieving successful expression of multiple constructs within a single cell is less likely compared to single constructs, influenced by various probability factors, including uptake and successful expression. **b)** Fluorescence microscopy images after the transfection of primary human T cells on 400 nm with three mRNA constructs (genes: GFP, mCherry, Firefly Luciferase) (scale bar, 5 μm). **c)** Flow cytometry data (GFP and mCherry channels) following multi-mRNA transfection on both porous (400 nm) and non-porous substrates. **d)** Analysis of the flow cytometry data and categorizing cells according to the number of expressed genes, with single, double and triple mRNA expression. Statistical significance was evaluated through a two-way ANOVA test (*: p < 0.05, **: p < 0.01, ***: p < 0.001).

#### CAR-T cells

We assessed the efficiency of delivering lentiviruses to generate CAR-T cells from primary human T cells. Compared to the expression of GFP constructs, the expression of CARs is more complex. The complexity is most likely related to presence of multiple components in CARs, including antigen recognition domains, signaling domains, and other functional elements.^24^ In order to produce CAR-T cells, primary human T cells were activated with αCD3/CD28 antibodies 24 hours before exposure to lentiviruses. Subsequently, T cells were cultured on both porous (400 nm) and non-porous surfaces, and the lentiviruses carrying anti-CD19 CAR constructs with a GFP marker (CAR19-GFP) were introduced to the culture media with no additives or enhancers. On Day 4, flow cytometry analysis revealed that the nanoporous platform significantly enhanced the delivery of CAR19 constructs in primary human T cells (52.5%) compared to flat surfaces (30.8%), without compromising cell viability. Notably, the increased expression level of GFP (MFI) might also suggest a higher rate of successful integration (Figure 4).

**Figure 4.**
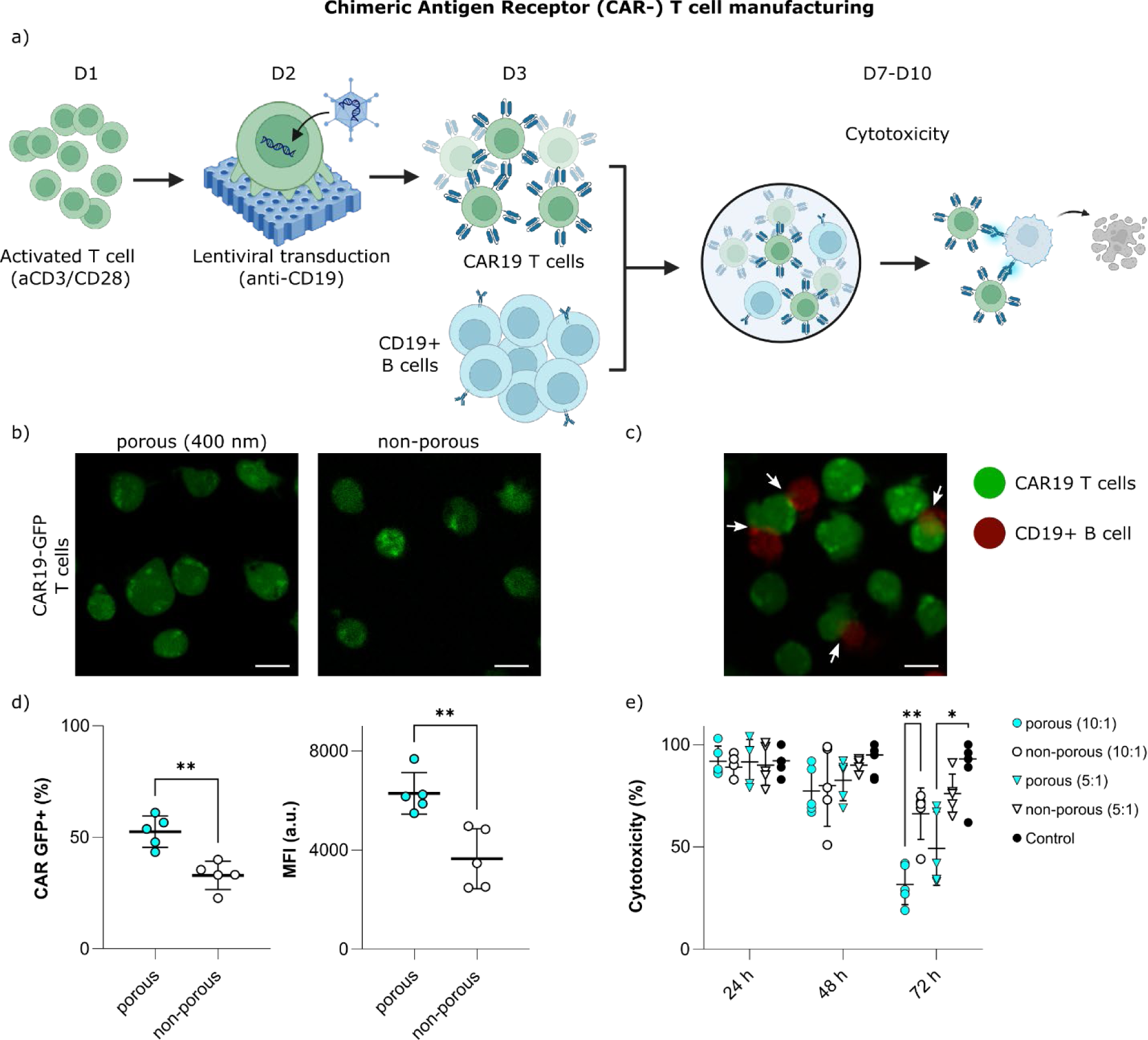
CAR T cell manufacturing. **a)** A schematic depiction of the experimental procedure: culturing antibody-activated primary human T cells (aCD3/CD28) on porous (or non-porous) surfaces for CAR lentiviral transduction, and the subsequent cytotoxicity assay by co-culturing CAR T with B cells (CD19+). **b)** Fluorescence microscopy images after the transduction of primary human T cells with CAR19-GFP lentiviruses for porous (400 nm) and non-porous conditions (scale bar, 5 μm). **c)** Fluorescence microscopy images of CAR19-GFP T cells (green) and B cells (red) co-cultured, showcasing the formation of synapses between the two cell types (scale bar, 5 μm). **d)** Analysis of flow cytometry data showing percentage of the transduced CAR19 T cells (GFP+), and the mean fluorescence intensity of the GFP+ cells under porous (400 nm) and non-porous conditions. **e)** Analysis fluorescence measurements of mCherry-expressing B cells co-cultured with varying ratios of CAR19 T cells (10:1 or 5:1), illustrating the cytotoxicity of CAR19 T cells (on porous 400 nm vs non-porous surfaces) over a 72-hour period. The control experiment involved culturing non-transduced T cells with B cells at a 10:1 ratio.

We evaluated the anti-cancer effectiveness of CAR19-T cells in an in vitro setting (Figure 4e). CD19-expressing Ramos B cells were chosen as a model for blood cancer (Lymphoma). To assess suppression of cancerous B cells by CAR19-T cells, we co-cultured the two cell types for a duration of 72 hours. B cells were transfected to produce fluorescent signal (mCherry), which was used as a readout for measuring the killing rate using a fluorescence plate reader. CAR19-T cells were co-cultured with B cells on Day 7 and at different ratios (10:1 and 5:1 ratio for T:B cells). Activated T cells (without transduction) were co-cultured with Ramos cells as a control experiment. Optical microscope images show the capability of CAR19-T cells in targeting B cells (CD19+) by forming close contacts (Figure 4b). CAR19-T cells from both porous and non-porous surfaces demonstrated suppression of the mCherry signal after 72 hours in comparison with the control sample, indicating the cytotoxicity of T cells. In general, CAR19-T cells produced on porous surfaces had a higher cytotoxicity (up to 80%) towards Ramos cells compared to cells from non-porous surfaces (up to 30%). A significant reduction in B cell viability was observed with the 10:1 cell ratio compared to 5:1 in porous surfaces. On non-porous surfaces, 5:1 cell ratio did not result in any significant changes in B cell viability. These results highlight the potential of nanoporous surfaces for efficient transduction of therapeutic agents into primary human T cells, with potential application in immunotherapy.

### On the mechanism of enhanced uptake

The enhanced cellular uptake on nanoporous surfaces can be attributed to factors such as increased cell contact with the surface and the stimulation of membrane ruffling on the cell membrane.^16,25^ These factors, in turn, can promote a range of uptake mechanisms, such as receptor-mediated and non-receptor-mediated endocytosis.^26,27^ Receptor-mediated endocytosis, such as clathrin- or dynamin-mediated endocytosis, is a typical cellular pathway for the entry of viral vectors and other biologics, allowing selective regulation of the uptake of substances that bind to membrane receptors.^27^ However, the observation that a variety of vectors (e.g. lentiviruses, baculoviruses, mRNA) and both small and large molecules exhibit increased cellular uptake on nanoporous surfaces suggests a mechanism that differs from receptor-mediated entry routes. We hypothesize that uptake of exogenous materials by cells on nanoporous surfaces is predominantly driven by fluid-phase endocytosis, particularly macropinocytosis.^28^

Macropinocytosis is a form of fluid-phase endocytosis used by cells to engulf volumes of extracellular fluid and its contents.^27^ During this process, the cell extends its membrane to form ruffles or protrusions on the cell surface (macropinosomes), which capture and transport external fluid along with its contents into the cell. Macropinocytosis takes place in a wide range of mammalian cells such as macrophages, dendritic cells, neutrophils, fibroblasts, epithelial cells, and T cells. ^29–31^ It is known to be a highly efficient mechanism in immune cells for capturing pathogens and antigens, contributing to immune responses.

Macropinocytosis is an actin-dependent endocytic process and it has been shown that chemicals that lead to an increased formation of membrane ruffles (such as PMA, growth factors, and modified LDLs), also lead to efficient uptake of exogenous materials.^29,32^ Nanoporous surfaces stimulate membrane ruffling in a manner similar to chemical inducers, but by employing physical cues.^16,33,34^ The surface’s topographical features prompt cells to react, by extending actin-rich membrane protrusions within the pores. The membrane protrusions are dynamic in nature and their formation/retraction is linked with cell crawling on the surface. These dynamic protrusions are continuously in motion, allowing them to capture and engulf volumes of extracellular fluid and its contents, making it a highly efficient uptake process. The outcome of this process is the generation of vesicles, including micropinosomes (typically less than 100 nm in diameter) or macropinosomes (ranging from 200 nm to 1 μm in diameter), which in part is determined by the shape and thickness of actin-rich protrusions.^35^

To determine if membrane protrusion induced by nanotopography can engage in macropinocytosis, we tested the ability of mouse pre-osteoblasts to endocytose 10 kDa and 70 kDa rhodamine B-dextran (Rhod-Dex), that are used commonly as macropinocytosis probes.^27,31^ Dextrans are globular proteins, with approximate Stoke’s diameters of 4.7 nm and 11.6 nm, for 10 kDa and 70 kDa molecules, respectively.^31^ Due to its large size, 70 kDa Rhod-Dex provides more specificity for macropinosomes, as its uptake is restricted to fluid phase uptake. For 10 kDa Rhod-Dex, diffusion through the cell membrane enables an additional route for cellular entry.^27,31^ As assessed by fluorescence measurement, mouse pre-osteoblasts readily took up both 10 kDa and 70 kDa Rhod-Dex on both porous (400 nm) and non-porous surfaces (Figure 5 and **Figure S7**).

**Figure 5.**
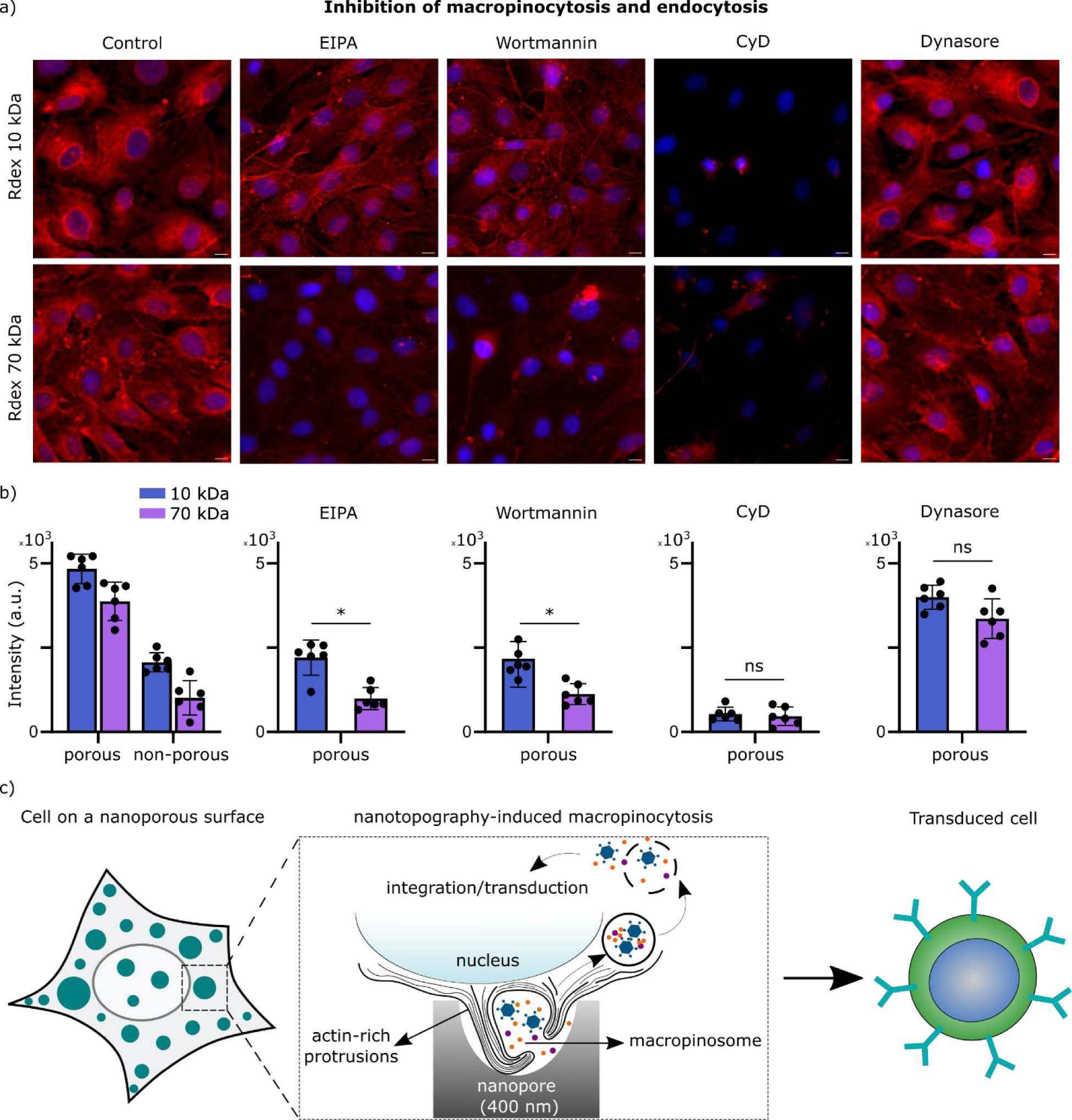
Mechanism of enhanced uptake on porous surfaces. **a)** Confocal fluorescence microscopy images showing uptake of Rhodamine B-Dextran (10 or 70 kDa) by mouse pre-osteoblast cells cultured on porous (400 nm) PCL films (scale bar, 5 μm). The nucleus was stained with Hoechst. Cells were pre-treated with various inhibitors (EIPA, Wortmannin, CyD, and Dynasore) before incubation with Rhodamine B-Dextran molecules. **b)** Fluorescence intensity measurement of Rhodamine B-Dextran uptake by the cells using a plate reader, with inhibition by the respective inhibitors. **c)** A schematic representation of nanotopography-induced micropinosome formation and downstream cellular processes leading to cell transduction.

We employed specific inhibitors to verify that macropinocytosis is responsible for the uptake of both small and large molecular weight probes, allowing us to differentiate macropinocytosis from other endocytic processes. These probes include, 5-(N-Ethyl-N-isopropyl) amiloride (EIPA), Cytochalasin D (CyD), Dynasore and Wortmannin. EIPA is a specific and effective probe to inhibit formation of macropinosomes by blocking a plasma membrane Na+/H+ exchangers.^27,31^ CyD disrupts actin polymerization, crucial for actin-rich structures in macropinocytosis.^16^ Dynasore is a cell-permeable molecule that inhibits dynamin-dependent endocytosis through blocking the dynamin GTPases.^36^ Wortmannin is an inhibitor of phosphoinositide 3-kinase (PI3K), which is essential for the cell surface ruffling associated with macropinocytosis.

Using both porous (400 nm) and non-porous surfaces, we observed that EIPA largely reduced uptake of both Rhod-Dex molecules (10 kDa and 70 kDa), especially for 70 kDa Rhod-Dex on porous surfaces. Stronger dependence of 70 kDa Rhod-Dex on macropinosome blocker suggests that macropinocytosis is the predominant mechanism on nanoporous surfaces.

Addition of CyD to the cells also reduced cellular uptake in a concentration-dependent fashion. The notable 90% decrease in cellular uptake on porous surfaces at low CyD concentrations (10 μM) highlights the important role of actin-rich protrusions in the uptake process. The impact of CyD addition on the uptake of 70 kDa Rhod-Dex was more pronounced than that of 10 kDa, exhibiting a steeper slope in the uptake-concentration curve (**Figure S8**).

Blocking dynamin-mediated endocytosis using Dynasore (100 mM) did not lead to a significant change in the uptake of 10 kDa and 70 kDa Rhod-Dex on nanoporous surfaces, however, on non-porous surfaces the uptake of 10 kDa Rhod-Dex was reduced (**Figure S7**). Transferrin-AF647 was used as a positive control to validate the influence of Dynasore treatment, which is a molecule that is internalized through receptor-mediated endocytosis. The results show that the uptake of transferrin was reduced by Dynasore treatment (**Figure S9**). Therefore, it is concluded that the uptake of Rhod-Dex molecules is dynamin-independent and it is a fluid-phase endocytosis.

Inhibition of PI3K using wortmannin (200 nM) significantly reduced the internalization of 10 and 70 kDa Rhod-Dex on both porous and non-porous surfaces. Similar to the impact observed with EIPA treatment, a more noticeable impact was observed for the larger 70 kDa Rhod-Dex molecules on porous surfaces.

Temperature also plays a significant role in influencing macropinocytosis, which demands substantial cellular energy. When cells were incubated at 4 °C, a significant reduction in cellular uptake was observed for both 10 and 70 kDa Rhod-Dex molecules. This reduction was observed regardless of whether the cells were cultured on porous or non-porous surfaces. There was no significant difference between the cellular uptake on porous and non-porous culturing conditions at 4 °C (**Figure S10**).

Moreover, macropinocytosis is a rapid and dynamic process compared to other endocytic mechanisms. As shown earlier (Figure 2, Figure S5), the rate of uptake on porous surfaces was considerably faster. Within the first hour of incubation with siRNA, GFP decay rate on porous surfaces (400 nm) reached ∼0.3 1/sec, which was an order of magnitude higher than the decay rate on non-porous surfaces (Figure S5). This suggests that the cellular mechanism underlying the uptake on nanoporous surfaces must be significantly faster than receptor-mediated endocytosis, which further corroborates the dominance of fluid-phase uptake on porous surfaces.

As shown by previous studies, macropinosomes typically formed at the plasma membrane have a diameter larger than 200 nm.^37,38^ This implies that in order for macropinosomes to be created within a material’s pores, there must be adequate space available to allow the actin-rich protrusions to generate circular ruffles of larger 200 nm in size. This is possible only when the material’s pore size exceeds 200 nm. Our observations are in line with the size for macropinocytosis, as no significant difference in cellular uptake was observed when comparing 100 nm pores to non-porous surfaces (Figure 2). However, a notably higher uptake was observed on 400 nm pores compared to 200 nm and 1000 nm pores (Figure 2). This observation could be due to the fact that 400 nm pores provide an optimal size for actin-rich protrusions to efficiently form circular ruffles. The 200 nm pores might limit the process, possibly because they restrict the formation of circular ruffles due to tight spatial confinement.^16^ On the other hand, the 1000 nm pores may not create the ideal conditions for efficient macropinosome formation, possibly due to the thicker actin-rich protrusions in larger pores compared to typical macropinosomes. Larger pores lead to actin-rich structures that are more stable and substantial, which could limit the closure of the protrusions through membrane fission. This issue is particularly notable as the 1 µm protrusions in T cells make up a relatively substantial portion, accounting for approximately 10-20% of the cell’s size, which could be impractically large for macropinosome formation.

The shape of pores, whether open or blind, appears to have varying effects on this process as well (Figure S2). The rounded-bottom structure of blind pores likely facilitates the engulfing of extracellular material into these circular ruffles, which are then internalized as macropinosomes. In contrast, straight pores may not provide the same conducive environment for the circular formation and engulfing process, which is why they may contribute differently to the overall process of macropinocytosis. Actin protrusions are typically oriented and guided along the direction of the channel, making it challenging for them to form a U-turn and to encapsulate extracellular material in the same manner as circular ruffles. This limitation in the geometry of straight channels could hinder or slow down the process of macropinocytosis in such environments.

Also electrostatic interactions play an important role in endocytosis pathways, excluding the fluid-phase uptake. The negligible impact of chemical enhancers, such as polybrene (Figure S4), on porous surfaces in our experiments provides compelling evidence against the involvement of electrostatic charges in the uptake process on porous surfaces, further supporting the macropinocytosis as the major mechanism for enhanced cellular uptake, as electrostatic charges do not play a role in fluid-phase uptake.

Moreover, primary human T cells actively utilize macropinocytosis to uptake extracellular amino acids into their endolysosomal compartments, promoting T cell growth and metabolism. The expression of the Rapamycin complex (*mTORC*) gene, crucial for promoting T cell growth and metabolism, is closely associated with macropinocytosis.^30^ We measured early expression of *mTORC* (6 hours) in non-activated primary human T cells cultured on porous (200 nm and 400 nm) and non-porous surfaces. *mTORC* was upregulated on porous surfaces compared to non-porous surfaces, with a significantly higher expression observed on the 400 nm porous structure (**Figure S11**).

Furthermore, we investigated the regulation of macropinocytosis on nanoporous surfaces, based on existing literature.^16^ The transcriptomics data obtained from non-activated primary human T cells that were cultured on porous (200 nm) and non-porous membranes was used to study the regulation of genes encoding for macropinocytosis induced by nanotopography. As shown by the heatmap in **Figure S12,** nanoporous surfaces trigger both upregulation and downregulation of genes associated with macropinocytosis (according to gene ontology analysis, GO:0044351). Most notably, is the upregulation of genes involved in actin nucleation and branching (*CDC42*, *RHOA*, *WASH*), and downregulation of genes involved in receptor-mediated endocytosis (*DNM2*, *APPL1*, *RAB5*) on nanoporous surfaces. This analysis further corroborates with enhanced fluid-phase uptake on nanoporous surfaces and suppressed receptor-mediated endocytosis.

In summary the enhanced uptake mechanism of cells cultured on nanoporous surfaces can be described as follows: The mechanical deformation of the cell membrane during cell movement on nanoporous surfaces results in transient membrane protrusions within the scale of the pore size, including macropinosomes. Blind pores facilitate the necessary membrane remodeling for macropinosomes formation, ultimately increasing the uptake of fluids and exogenous materials through macropinocytosis. Macropinocytosis stands out due to its efficiency in taking up large volumes of extracellular material and its ability to bypass endosomal degradation, increasing the likelihood of intact cargo reaching the cytoplasm and the nucleus.

## Conclusion

In conclusion, this study explored the impact of nanotopographical cues from nanoporous surfaces on the cellular uptake of various biomolecules. The results revealed a significant increase in cellular uptake efficiency on nanoporous surfaces, with the process being size and morphology-dependent, reaching optimal efficacy with blind pores of 400 nm in diameter. The enhanced genetic transduction observed encompasses various vectors, such as lentiviruses, baculoviruses, mRNA molecules. Other small or large molecules such as polysaccharide (Dextran) or antibodies also have an easier route of entry to the cells on nanoporous surfaces. The applicability of the introduced method in therapeutic and biotechnological applications was demonstrated by effectively delivering multiple mRNA constructs into individual primary human T cells, as well as by the efficient production of potent CAR-T cells.

The approach for enhancing cellular uptake on nanoporous surfaces is not limited by a cargo-specific entry mechanism, and it provides a generic path for the entry of different cargoes or viruses. The independence from cargo properties and cell characteristics establishes it as a versatile and widely applicable technique, demonstrated by the efficacy of the method in both adherent and non-adherent cells, as well as primary immune cells.

Furthermore, our data analysis including inhibitor studies targeting different uptake pathways, pointed towards macropinocytosis as the primary mechanism responsible for the improved cellular uptake and transduction. Macropinocytosis is a broadly applicable nonspecific uptake mechanism, which is fast and efficient, bypassing endosomal degradation and minimizing immunogenicity in contrast to artificial delivery approaches. These findings suggest that nanoporous surfaces can guide cells to achieve improved uptake, providing applications in areas such as gene delivery and other biomedical processes related to cell engineering.

## Methods

### Nanoporous surfaces

Nanoporous polycaprolactone (PCL) membranes were fabricated through casting as reported elsewhere. Briefly, the PCL polymer with an average molecular weight of 80,000 g/mol (Sigma-Aldrich, 440744) was mixed with dichloromethane (DCM; fisher scientific, 10127611) and 0.1% Tween. Air bubbles were introduced into the mixture through sonication at 45 kHz. The size of air bubbles undergoes variations during sonication. Solutions with PCL concentrations of 5-10% were sonicated for durations spanning from 30 minutes to 300 minutes, resulting in the formation of air bubbles within the range of 100 nm to 1000 nm, that contribute to the creation of nanopores within the PCL structure. The solution was poured to the bottom of the well plates until dried. The surface of the polymer film was sterilized using 70% ethanol for 10 min, and washed with PBS immediately before culturing the cells.

### Cell culture

#### Mouse pre-osteoblasts

MC3T3-E1 cells were grown in α-MEM culture media without ascorbic acid (Gibco, A1049001, Thermo Fisher Scientific), and were supplemented with 10% Fetal Bovine Serum (FCS) (Gibco, fisher scientific, 11560636) and 1% penicillin/streptomycin (Sigma-Aldrich, P4333), at 37°C with 5% CO2. The cells were detached from culture flasks, using TrypLE (Gibco, fisher scientific, 11528856) for 5 minutes at 37°C, and were resuspended to a concentration of 2×10^5^ cells/mL, prior to culturing on nanoporous surfaces. A volume of 200 μL was used for culturing cells in 96-well plates, for 4-24 hours prior to uptake analysis.

#### Primary human T cell isolation

Human peripheral blood mononuclear cells (PBMCs) were isolated using Ficoll-Paque (GE Healthcare Life Science) from buffy coats donated by healthy, anonymous individuals, sourced from the Blood Centre at Akademiska Sjukhuset (Uppsala, Sweden). CD4+ and CD8+ T cells were positively sorted from the PBMCs utilizing the StraightFrom® Buffy Coat CD4/CD8 MicroBead (Miltenyi Biotec) and subsequently separated using the QuadroMACS™ Separator (Miltenyi Biotec).

### T cell activation

T cells were activated using αCD3/CD28 beads (T Cell TransActTM; Miltenyi Biotec), according to the manufacturer’s protocol. T cells were cultured in TexMACS GMP Medium (Miltenyi Biotec), which was supplemented with IL7 (155 U/mL), IL15 (290 U/mL), and 1% penicillin/streptomycin. The cells were incubated at 37 °C in a humidified 5% CO2 incubator, and the cell concentration was maintained at 10^6^ cells/mL. Activated T cells were cultured on porous or non-porous surfaces 24 hours after activation.

### Ethical considerations

In this study, anonymized human PBMCs were collected from donors, which did not require ethical approval. No alternative materials were used that demand ethical considerations.

### Cellular uptake

Various constructs, including lentiviruses, baculoviruses, mRNAs and antibodies were tested for their cellular uptake efficiency.

#### Lentiviruses

In-house preparation was carried out for GFP lentiviruses (see below). 5 μL of the stock solution was added to the pre-osteoblasts 24 hours after seeding. The cells were cultured with the virus particles at 37 °C in 5% CO2 for 24 hours. The cells were rinsed twice with warm PBS, and the intensity of GFP fluorescence was quantified using a plate reader (Tecan Infinite 200) at an excitation/emission wavelength of 490/520 nm. The experiments were conducted with 6 replicates.

For T cell studies, primary human T cells were employed to assess transduction efficiency. T cells were seeded at a concentration of 10^6^ cells/mL. Subsequently, 500 μL of activated or non-activated cells from three distinct donors were cultured on surfaces, both porous and non-porous, placed in 24-well plates. The experiments were conducted with at least 5 replicates. Cells were incubated with the viruses for 24 hours. Flow cytometry (CytoFLEX S, Beckman Coulter) was utilized for primary human T cells to assess GFP expression and determine the percentage of cells, as well as mean fluorescent intensity of GFP per cell, that underwent transduction.

#### Baculoviruses

GFP baculoviruses were obtained from Invitrogen (Invitrogen, B10383), with a stock solution concentration of 10^8^ viral particles/mL. 2 μL of the stock solution was added to the pre-osteoblasts 24 hours after seeding, and the optical measurement was performed similar to lentivirus uptake measurements.

#### mRNAs

StemMACS™ Nuclear eGFP mRNA were used for mRNA uptake analysis, which encodes an enhanced green fluorescent protein (eGFP) inside of the nucleus. The concentrated stock solution had a concentration of 100 ng/μL. 10 ng of the solution was added to the pre-osteoblast cells and were incubated for 24 hours. The cells were rinsed and the signal was measured using the plate reader.

#### Antibodies

Antibodies uptake was determined by measuring the fluorescent intensity of a fluorescently-tagged antibody (Alexa Fluor® 647 anti-Tubulin-α, 0.5 mg/mL, BioLegend 627908) at a dilution ratio of 1:2000. Optical intensity measurements were conducted 24 hours after incubation with the antibody at an excitation/emission wavelength of 650/670 nm.

### Cell viability measurements

Cell viability was measured for pre-osteoblast cells that were incubated with lentiviruses, baculoviruses or mRNAs. LIVE/DEAD staining assay (Invitrogen™ L3224, Thermo Fisher Scientific) was used according to the manufacturer protocol. 400 nm porous PCL were used as nanoporous surfaces, whereas cells cultured on non-porous surfaces served as the control for cytotoxicity assessment. Cytotoxicity was determined by comparing the relative viability of the cells with that of the non-porous samples.

### GFP lentivirus production

The lentiviral plasmid carrying a GFP construct under the CMV promoter (pBMN-CMV-GFP) was employed to evaluate transduction efficiency via flow cytometry. Plasmids for lentivirus production were generated by expanding a bacterial strain containing the lentiviral plasmid and helper plasmids (pLP1, pLP2, and pLP-VSVG) over two days. Subsequently, plasmid extraction was carried out using the Plasmid Plus Midi Kit (QIAGEN®) following the manufacturer’s protocol. The integrity of both the GFP plasmid and the helper plasmids was confirmed through gel electrophoresis and imaging using GelDocTMXR+ (BioRad).

For the production of lentiviral vectors (LV(GFP)), human embryonic kidney 293T (HEK293T) cells were cultured in 20 mL of DMEM glutamax growth medium (containing 10% FBS, 1% sodium pyruvate, 1% penicillin/streptomycin, and geneticin at 500 μg/μL, Gibco) at 37°C in a 5% CO2 environment. Cells were transfected with the lentiviral plasmid (CMV-GFP; 600 ng/mL), pLP1 (300 ng/mL), pLP2 (300 ng/mL), and pLP/VSVG (300 ng/mL) for 24 hours. Virus-containing supernatants were collected at 48 hours and 72 hours after transfection and concentrated using ultracentrifugation (Ultraclear, Beckman 344058). The transduction capacity of the viral vector product was confirmed by culturing non-transfected HEK293T cells with the virus after 24 hours and assessing via fluorescent imaging.

### CAR19-GFP lentivirus production

The human CD19-targeting Chimeric antigen receptor (CAR) sequence, which contains single chain fragment derived from FMC63 clone, the human CD3zeta and CD137 signalling domains as described previously.^12^ The encoding sequence was cloned into a third-generation self-inactivating (SIN) lentiviral vector under the control of the elongation factor-1 alpha (EF1a) promoter. A green fluorescent protein was incorporated after the CAR cassette and separated by a self-cleaving T2A sequence. The lentiviral vector was designated LV(CAR19-GFP).

Vesicular stomatitis virus (VSV)-G pseudotyped lentiviral particles were produced in HEK-293T cells. Cells were transfected with the lentiviral vector plasmid and the pLP1, pLP2 and pLP/VSVG (Invitrogen) packaging plasmids at a 1:1:1:2 molar ratio using polyethyleneimine (Sigma-Aldrich). Viral supernatant was collected 48 and 72 h post transfection, filtered (0.45 μm), and concentrated by ultracentrifugation at 75,000 × g for 90 min at 4 °C. The viral pellet was resuspended in DMEM and stored at −70 °C until further use.

### CAR-T cell manufacturing

T cells were first activated with αCD3/CD28 antibodies 24 hours before lentivirus exposure. Activated T cells were cultured on porous and non-porous surfaces in 48 well plates, and the lentiviruses (CAR19-GFP) were added immediately (15 μL of virus per 250 mL of cell suspension; 10^6^ cells/mL). The cells were kept in an incubator at 37 °C with 5% CO2, and the culture media was exchanged at least every two days. The transduction efficiency of the modified T cells was assessed using flow cytometry three days after the transduction process (Day 4 of activation). For flow cytometry, T cells were collected from the surfaces and washed with PBS, followed by staining with a viability dye (Thermo Fisher Scientific, P3566) for 20 min at 4 °C. The cells were washed and GFP expression was measured immediately with a CytoFLEX S flow cytometer (Beckman Coulter) and data was processed using Cytoflow software.

### CAR-T cell cytotoxicity

Cytotoxicity assessment was conducted using CAR-T cells on Day 7 after activation. Co-culturing CAR-T cells with mCherry expressing Ramos B cells was performed to measure killing rates using a fluorescence plate reader. T and B cells were both resuspended in RPMI culture media (Gibco, RPMI 1640 Medium, GlutaMAX™ Supplement, 61870036) supplemented with 10% FCS and 1% penicillin/streptomycin, at a concentration of 10^4^ -10^5^ cells/mL. The two cell suspensions were mixed in 5:1 and 10:1 ratio and a final volume 200 μL was added to each well of a 96-well plate. The cells were cultured at 37 °C with 5% CO2 for durations of 24, 48 and 72 hours. At each interval, the culture media were replaced with PBS, and the fluorescent signal was measured using a plate reader (Tecan, Infinite 200).

### Multiple mRNA transfection

T cells were cultured on 400 nm porous surfaces and non-porous surfaces (20,000 cells per well). mRNA Cap1-mCherry, mRNA Cap1-FireflyLuciferase and mRNA Cap1-GFP were purchased from PackGene, USA. Transfection was carried out using Lipofectamine™ 3000 Transfection Reagent (Invitrogen, L3000001) according to the manufacturer’s protocol. Briefly, a total of 3 μg of mRNA (comprising 1 μg of each mRNA) was mixed with 100 μL of Opti-MEM medium (Gibco CTS Opti-MEM I Medium, 16386572). Lipofectamine™ 3000 Reagent was diluted in Opti-MEM in 6:100 ratio and it was mixed with the mRNA solution (1:1) for 10 min. 10 μL of the master mix was then added to the cells in each well of a 96-well plate. The cells were incubated 37 °C with 5% CO2 for 24 hours, and subsequently removed from the substrate and rinsed with PBS before measuring using a flow cytometer (CytoFLEX S flow cytometer, Beckman Coulter).

### Uptake characterization

#### Kinetics characterization

GFP expressing Jurkat cells were cultured on porous (200 nm and 400 nm pores) and non-porous surfaces 30 min before incubation with siRNA (Silencer™ GFP siRNA from Invitrogen, 10025764, 5 nmol). Negative Control siRNA from the kit was used as a negative control. A total of 40,000 cells were cultured in individual wells of a 96-well plate, with each well containing 200 μL of RPMI culture media. Subsequently, 2 μL of the siRNA stock solution (equivalent to 10 pmol) was added to each well. Prior to the 6-hour measurement course, the plate reader (Tecan Infinite 200) was pre-heated to 37 °C. During the measurement, data were collected at 4-minute intervals with excitation/emission wavelengths set at 490/520 nm. The experiments were conducted with 6 replicates for each condition.

To assess the impact of temperature on siRNA (5 nmol) uptake, the GFP expressing Jurkat cells were cultured on porous (400 nm) and non-porous surfaces, and were subjected to incubation at either 4 °C or 37 °C, for durations of 30 minutes, 1 hour, 2 hours, and 4 hours.

#### Pore size

To investigate the influence of pore size on cell uptake, pre-osteoblast cells were cultured on PCL films with different pore diameters. Cell culture conditions were as previous measurements (α-MEM without ascorbic acid 10% FCS and 1% pen/strep, at 37°C with 5% CO2, 2×10^5^ cells/mL). A volume of 200 μL was used for culturing cells in 96-well plates, for 4 hours prior to the addition of mRNA (StemMACS™ Nuclear eGFP mRNA). 10 ng of the solution was added to the cells and were incubated for 24 hours. The cells were rinsed and the signal was measured using the plate reader.

#### Pore shape

Whatman™ Polycarbonate Track-Etched Membranes as well as Whatman Anodisc inorganic filter membranes were used to investigate the pore shape effect [16]. Membranes with 200 nm and 400 nm pores were used to compare the transduction efficiency with round bottom PCL membranes. The membranes (13 mm) were placed in a 24-well plate and a volume of 400 μL of cells (2×10^5^ cells/mL) was added to each well and incubated for 4 hours at 37°C with 5% CO_2_. 20 ng of the nuclear-GFP mRNA was added to the cells and incubated for 24 hours. The cells were rinsed and detached from the surfaces using TrypleE, and were seeded on a 96-well plate with a total volume of 200 μL per well. The optical signal was measured using the plate reader 4 hours after seeding.

### Optical microscopy

For all fluorescence imaging, SP8 confocal laser scanning microscope (Leica) was used to acquire images using a 20× air objective, and Leica LAS X SP8 software. The nucleus was stained with Hoechst 33258 solution (1 mg/mL, Merck 94403) in 1:2000 dilution.

For high resolution imaging of actin protrusions, PCL films were coated on glass coverslips and cells were seeded on the surface (20,000 cells per sample) and incubated at 37 °C and 5% CO2 for 30 min. Cells were seeded on the surfaces (porous and non-porous, 50,000 cells per sample) and incubated for the planned duration. All the buffers were pre-warmed at 37 °C. Then the cells were pre-fixed for 1 min in 0.5% PFA (Formaldehyde, Polysciences, Inc) in PBS, followed by a fixation and permeabilization with 4% PFA and 0.1% Triton-X in PBS for 10 min. The samples were washed with PBS three times and incubated with 0.01% NaCl4 in PBS for 10 min to reduce the autofluorescence. The samples were then washed with PBS for three times and were incubated with AlexaFluor™ 488 phalloidin (Invitrogen, ActinGreen™ 488 ReadyProbes™ Reagent, 14879680) in 2% BSA in PBS for 90 min. The samples were washed with PBS three times before mounting with Prolong gold antifade reagent (Molecular Probes) on 1.5 H thick coverslips for imaging.

### Rhodamine uptake

Dextran-tagged Rhodamine-B (Rhod-Dex) with 10,000 Da and 70,000 Da molecular weights were from Invitrogen (Invitrogen 11466337 and 11590226). Mouse pre-osteoblast cells were cultured on 400 nm porous and non-porous surfaces in a 96-well plate 24 hours before the incubation with Rhod-Dex. Cellular uptake was quantified using a plate-reader (Tecan Infinite 200) with excitation/emission wavelengths set at 570/590 nm.

To assess the impact of temperature, pre-osteoblast cells were (w/o Dynasore treatment) subjected to incubation at either 4 °C or 37 °C, during the course of incubation with Rhod-Dex.

### Inhibition studies

The experiments were conducted in triplicates with two parallel biological replicates. All incubations were performed at 37 °C and 5% CO2. Pre-osteoblast cells cultured on porous (400 nm) and non-porous surfaces were pre-incubated for 30 minutes in either the absence (control) or presence of the inhibitors, after which the cells were washed with PBS and incubated with 1 mg/mL Rhod-Dex70 or Rhod-Dex10 for 1 hour. Subsequently the cells were washed with PBS.

5 mM amiloride (Invitrogen, 17197349) was used to inhibit the formation of macropinosomes. Cytochalasin D (CyD) (Sigma-Aldrich, C2618) was used to block actin polymerization at concentrations ranging from 0.1 to 100 µM. Dynasore (Invitrogen, 17220900) was used to inhibit dynamin-dependent endocytosis at a concentration of 10 µM. Wortmannin (fisher scientific, SE, 10706642) was used to inhibit PI3K (phosphoinositide 3-kinase) at a concentration of 200 nM. Transferrin-Alexa Fluor 647 Conjugate (Invitrogen, 11550766) at 1 mg/mL concentration was used to monitor receptor-mediated endocytosis, with and without Dynasore inhibition.

### RT-qPCR

Real-time quantitative polymerase chain reaction (RT-qPCR) was conducted on primary human T cells that were cultured on previously synthesized scaffold structures following a 6 hours seeding period. Total RNA extraction was carried out using the RNeasy Mini Kit from Qiagen (Qiagen, 74104). The isolated RNA was then assessed for quality with the 260/280 nm absorption ratio, ensuring a value greater than 2.0, using Nanodrop. To prepare cDNA, the iScript Advanced cDNA Synthesis Kit for RT-qPCR (1708891, Bio-Rad) was employed. Subsequently, RT-qPCR was performed utilizing SsoAdvanced Universal SYBR Green Supermix (Bio-Rad, 725270) within a CFX96 Connect RT PCR detection system (Bio-Rad). The reference gene for normalization of the acquired data was 18S RNA (RNA18S), and the primers were synthesized by Microsynth (Switzerland) at a concentration of 100 nmol/mL.

**Table 1.**
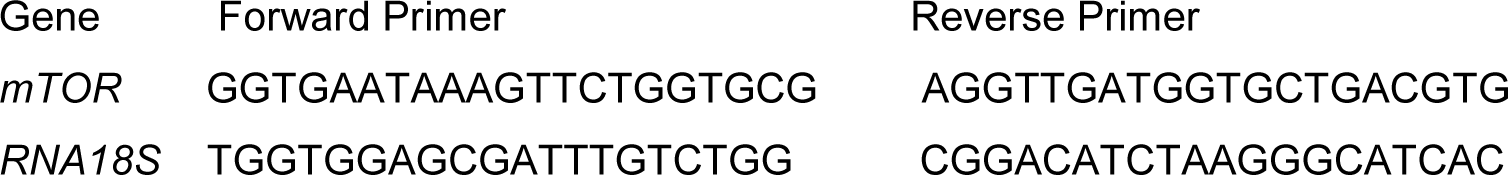
Primers sequences used for RT-qPCR:

### Gene transcriptomics analysis

Gene transcriptomics analysis was conducted using publicly available open-source data derived from primary human T cells cultured on both porous (200 nm) and non-porous surfaces (https://github.com/mortimus-p/T-cell-microvilli).

### Statistical analysis

All data were analyzed with GraphPad Prism. Normality of the distribution was assessed using the Shapiro-Wilk test, and group comparisons were conducted through a two-way ANOVA with a Tukey’s multiple comparisons test. The two factors encompassed experimental groups and biological replicates, with no observed significance between the biological replicates.

## Acknowledgement

We thank the funding support from Sweden’s Innovation Agency VINNOVA (Grant no: 2023-01495) and Carl Tryggers Stiftelse (CTS 22:2367). We thank our lab members Jing Ma, Lotte Duijn, Estefania Echeverri Correa and Vitalii Shtender for assistance during experiments. We thank Dirk Pacholsky and BioVis facility of Uppsala University for FlowCytometry. This work was conducted within the Additive Manufacturing for the Life Sciences Competence Center (AM4Life). The authors acknowledge financial support from Sweden’s Innovation Agency VINNOVA (Grant no: 2019-00029).

