## Supporting Information for "Nanotopography boosts cellular uptake by inducing macropinocytosis"

**Figure S1.** Confocal fluorescence microscopy images of mouse pre-osteoblast cells transduced with nuclear-GFP mRNA upon seeding on PCL films with 400 nm size, after 48 hours. There was a notable leakage of nuclear GFP into the cytosol which might be linked to the over expression of the proteins.

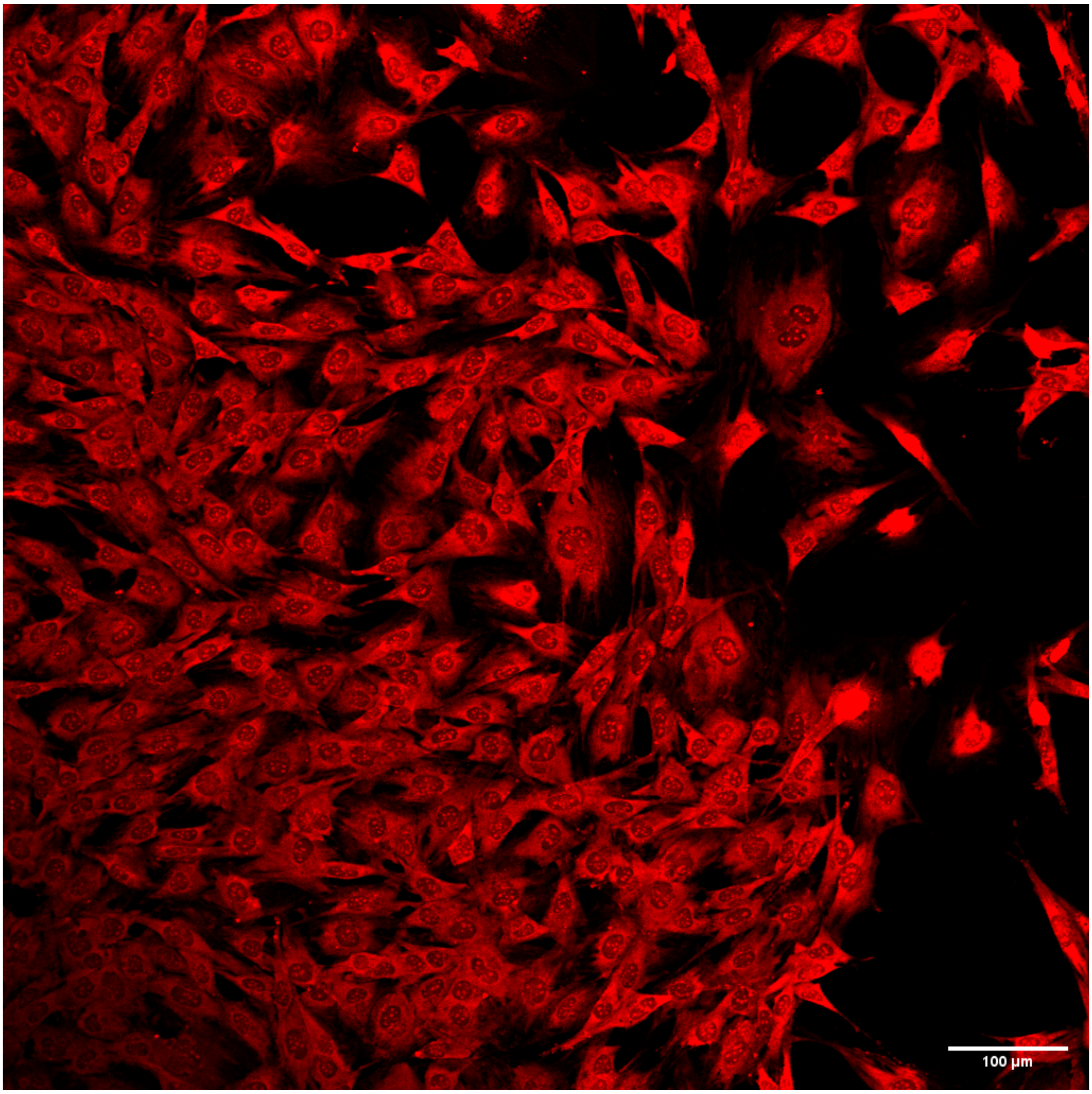

**Figure S2.** Influence of pore shape in cellular uptake. Mouse pre-osteoblast cells transduced with nuclear-GFP mRNA upon seeding on porous surfaces with different shape. AAO (anodic aluminum oxide) and Polycarbonate (PC) membranes had straight channels, while Polycaprolactone (PCL) film had blind pores. AAO films had a porosity of 40% and 50% for 200 and 400 nm pores, respectively. Porosity of ion track-etched PC membranes was 25%. Porosity of PCL films was 40%, measured by electron microscopy).
Quantitative uptake assessments were performed utilizing optical readings obtained with a plate reader. A scrambled mRNA (GFP-) was employed as the control. Statistical significance was determined through a two-way ANOVA test (*: $p < 0.05$, **: $p < 0.01$, ***: $p < 0.001$).

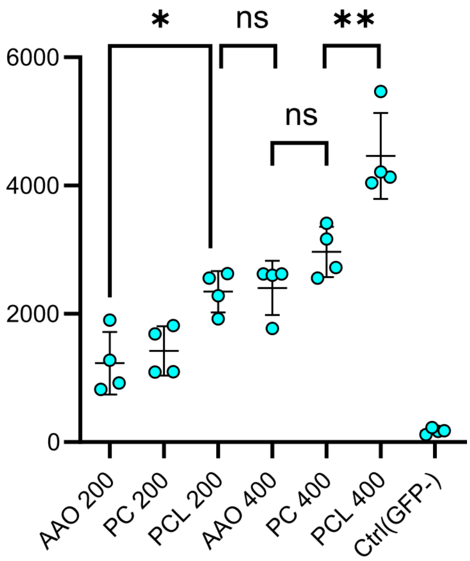

**Figure S3.** Scanning electron microscope image showing a cross-section of a pore in porous PCL film with an average pore size of 400 nm.

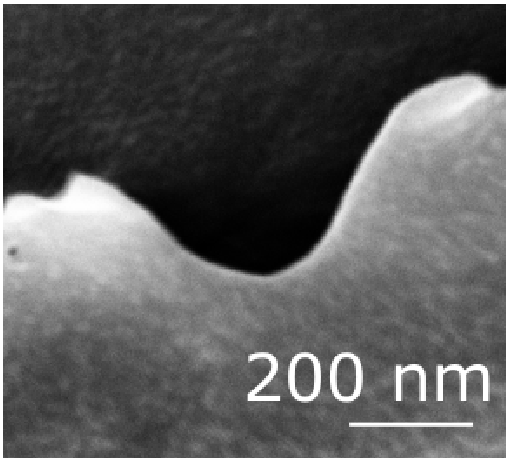

**Figure S4.** The influence of polybrene as a chemical enhancer in the uptake of GFP lentiviruses by primary human T cells. Polybrene, as a positively charged polymer, enhances lentiviral uptake through charge neutralization on the cell membrane and the viral envelope. The application of polybrene (10 µg/ml) resulted in a significant rise in cell transduction on non-porous surfaces, however, no significant effect was observed on porous surfaces.

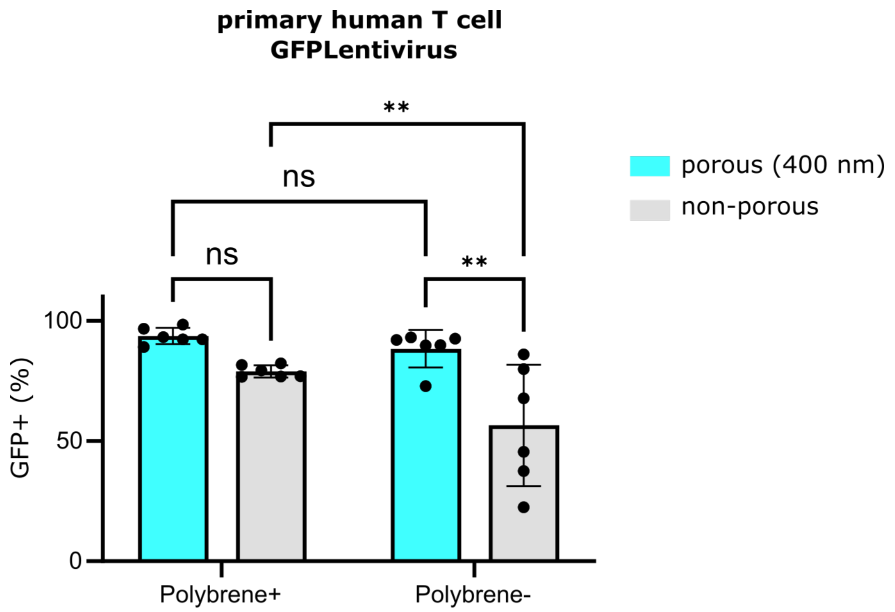

**Figure S5**. Decay rate of GFP signal (dI/dt). The derivatives are obtained from data presented in Figure 2c of the manuscript, where the uptake of siRNA by mouse pre-osteoblast cells was assessed on 400 nm, 200 nm, and non-porous films.

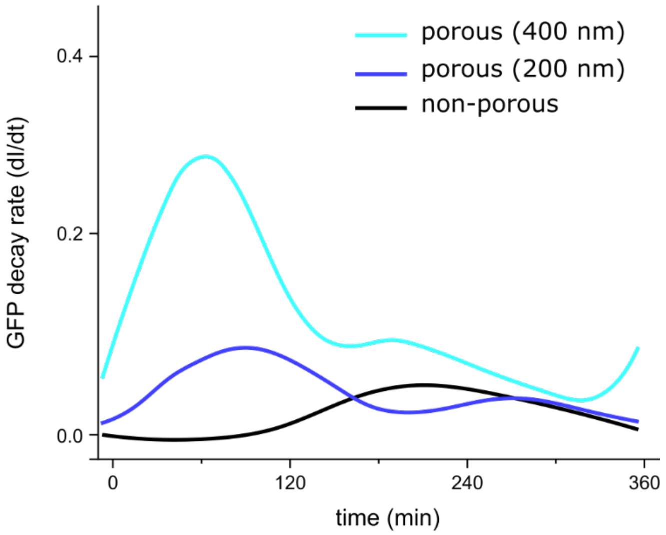

**Figure S6**. Influence of temperature on cellular uptake. Cells exposed to siRNA at 4 °C for durations of 30 minutes, 1 hour, 2 hours, and 4 hours showed minimal alteration in the GFP signal, observed consistently on both porous (400 nm) and non-porous surfaces.

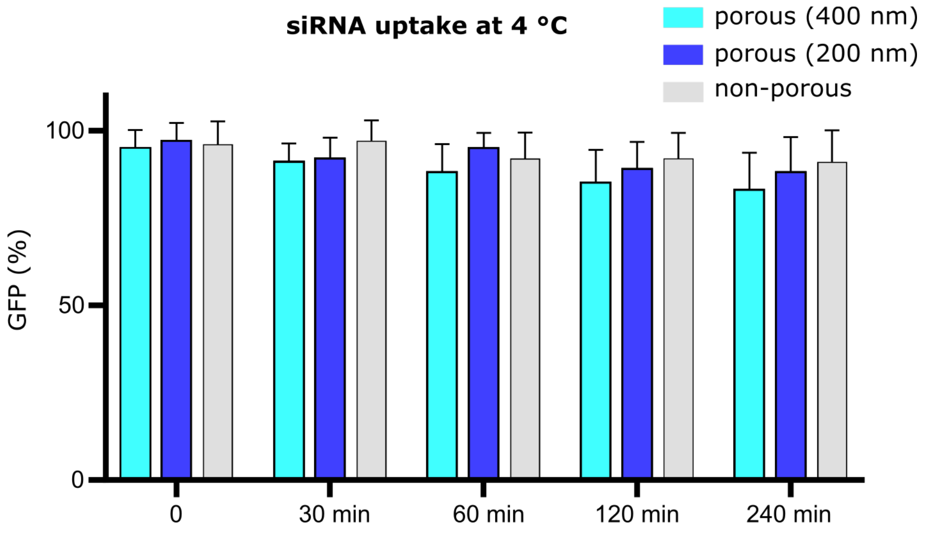

**Figure S7.** Inhibition of uptake on non-porous surfaces. Measurement of Rhodamine B-Dextran uptake by cells through fluorescence intensity using a plate reader, along with inhibition by the corresponding inhibitors.

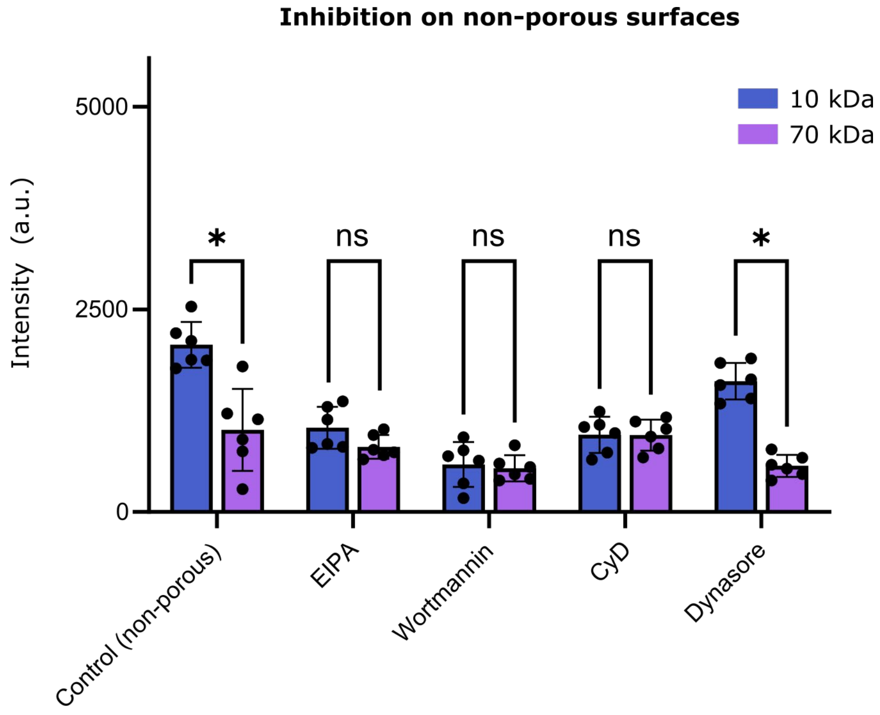

**Figure S8.** Influence of Cytochalasin D (CyD) treatment on uptake of Dextran molecules (10, 70 kDa). The introduction of CyD to the cells led to a concentration-dependent reduction in cellular uptake. The significant decline in cellular uptake on nanoporous surfaces at low CyD concentrations underscores the crucial involvement of actin-rich protrusions in the uptake process.

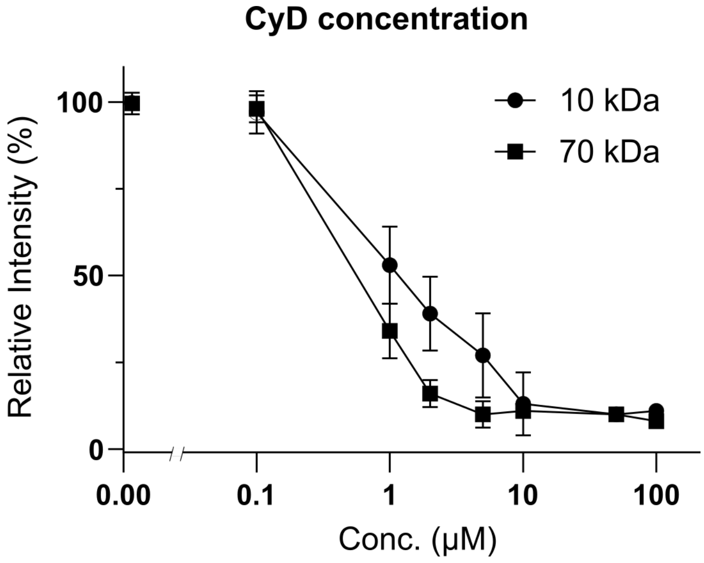

**Figure S9.** Transferrin-AF647 served as a control to confirm the impact of Dynasore treatment, a molecule internalized through receptor-mediated endocytosis. The results show a decrease in the uptake of transferrin upon treatment with Dynasore on both porous and non-porous surfaces.

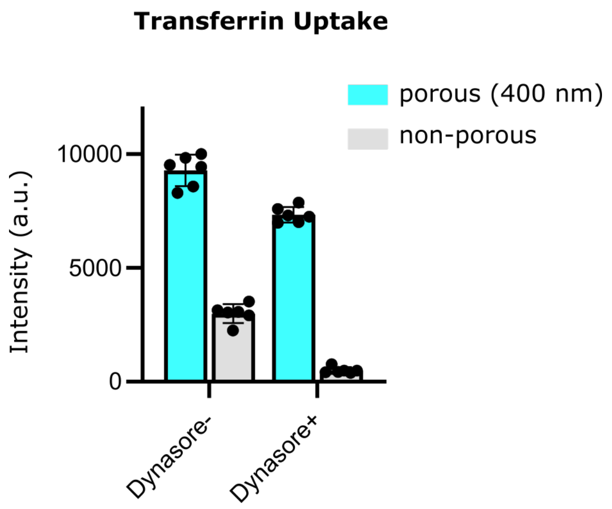

**Figure S10.** Influence of temperature in uptake of Dextran molecules (10 and 70 kDa). Incubating cells at 4 °C resulted in a decrease in cellular uptake for both 10 and 70 kDa Rhod-Dex molecules. This reduction was consistent regardless of whether the cells were cultured on porous or non-porous surfaces. No significant changes in cellular uptake was observed between porous and non-porous culturing conditions at 4 °C.

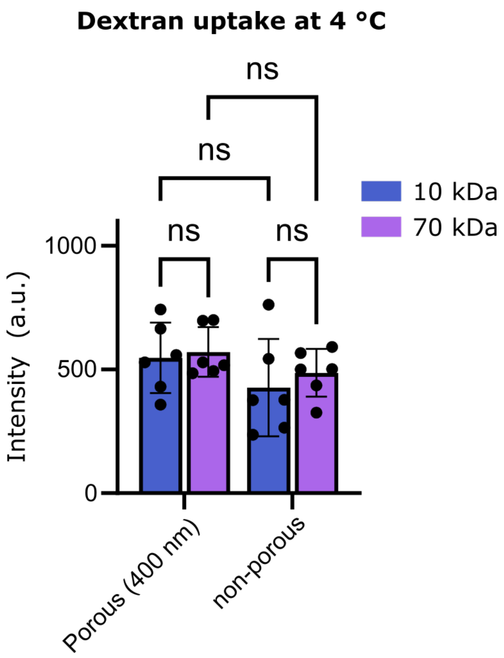

**Figure S11.** Relative RNA expression of *mTOR* by RT-qPCR. Primary human T cells were cultured on porous (200 and 400 nm) and non-porous surfaces for 6 hours. Fold expression is measured against the control sample (non-porous) and normalized by the reference gene RNA18S. The dash line shows the expression level in non-porous samples.

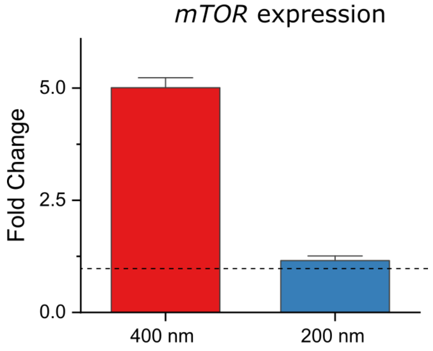

**Figure S12.** Transcriptomics analysis. Nano topography-induced alteration in the gene program of the primary human T cells after 4 h of seeding on the surfaces with 200 nm pore size (AAO surfaces). Nanoporous surfaces induce alterations in the expression of genes related to macropinocytosis (according to gene ontology analysis, GO:0044351). There is an increase in the expression of genes associated with actin nucleation and branching (*CDC42, RHOA, WASH*) and a decrease in the expression of genes linked to receptor-mediated endocytosis (*DNM2, APPL1, RAB5*) when cells interact with nanoporous surfaces.

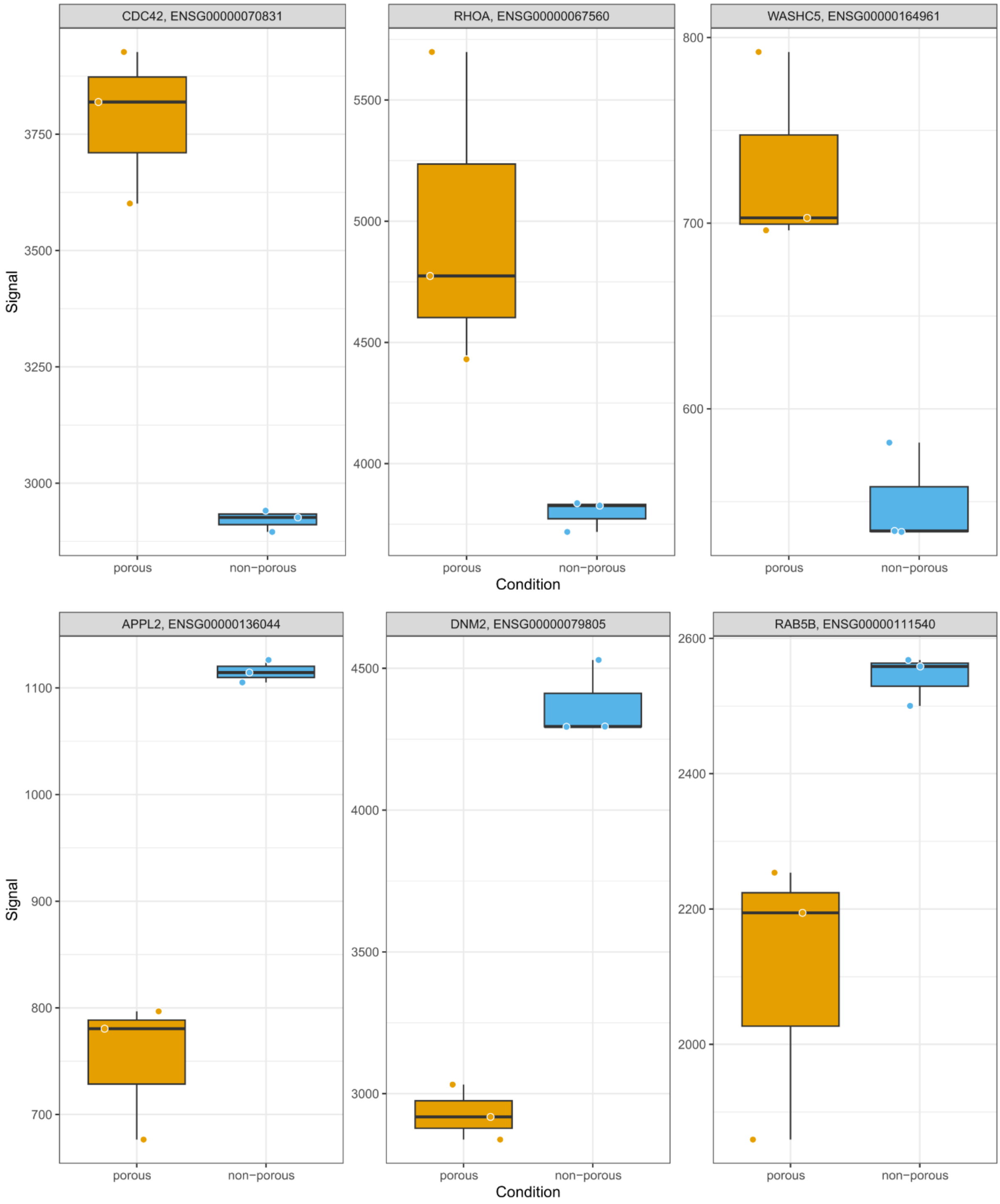
